# A blue-shifted genetically encoded Ca^2+^ indicator with enhanced two-photon absorption

**DOI:** 10.1101/2023.10.12.562058

**Authors:** Abhi Aggarwal, Smrithi Sunil, Imane Bendifallah, Michael Moon, Mikhail Drobizhev, Landon Zarowny, Jihong Zheng, Sheng-Yi Wu, Alexander W. Lohman, Alison G. Tebo, Valentina Emiliani, Kaspar Podgorski, Yi Shen, Robert E. Campbell

## Abstract

**Significance:** Genetically encoded calcium ion (Ca^2+^) indicators (GECIs) are powerful tools for monitoring intracellular Ca^2+^ concentration changes in living cells and model organisms. In particular, GECIs have found particular utility for monitoring the transient increase of Ca^2+^ concentration that is associated with the neuronal action potential. However, the palette of highly optimized GECIs for imaging of neuronal activity remains relatively limited. Expanding the selection of available GECIs to include new colors and distinct photophysical properties could create new opportunities for *in vitro* and *in vivo* fluorescence imaging of neuronal activity. In particular, blue-shifted variants of GECIs are expected to have enhanced two-photon brightness, which would facilitate multiphoton microscopy.

**Aim:** We describe the development and applications of T-GECO1 – a high-performance blue-shifted GECI based on the *Clavularia sp.*-derived mTFP1.

**Approach:** We used protein engineering and extensive directed evolution to develop T-GECO1. We characterize the purified protein and assess its performance *in vitro* using one-photon excitation in cultured rat hippocampal neurons, *in vivo* using one-photon excitation fiber photometry in mice, and *ex vivo* using two-photon Ca^2+^ imaging in hippocampal slices.

**Results:** The Ca^2+^-bound state of T-GECO1 has an excitation peak maximum of 468 nm, an emission peak maximum of 500 nm, an extinction coefficient of 49,300 M^-1^cm^-1^, a quantum yield of 0.83, and two-photon brightness approximately double that of EGFP. The Ca^2+^-dependent fluorescence increase is 15-fold and the apparent *K*_d_ for Ca^2+^ is 82 nM. With two-photon excitation conditions at 850 nm, T-GECO1 consistently enabled detection of action potentials with higher signal-to-noise (SNR) than a late generation GCaMP variant.

**Conclusion:** T-GECO1 is a high performance blue-shifted GECI that, under two-photon excitation conditions, provides advantages relative to late generation GCaMP variants.

## 1 Introduction

Genetically encodable calcium ion (Ca^2+^) indicators (GECIs), engineered from *Aequorea victoria* green fluorescent protein (avGFP),^1^ or its homologs, are powerful tools for enabling observation of intracellular Ca^2+^ dynamics. Among GECIs, the highly optimized jGCaMP series represents the tip of the spear with respect to pushing the limits of *in vivo* performance, particularly for the imaging of neural activity.^2–4^

The jGCaMP series has been iteratively and aggressively optimized for high sensitivity, high brightness under one-photon excitation, and fast kinetics, to great success.^4^ However, there are a variety of other desirable GECI properties that are unlikely to be realized with the avGFP-derived jGCaMP series, regardless of the extent of optimization. Such properties tend to be those that are intrinsic to the parent fluorescent protein (FP), such as higher two-photon brightness, fluorescence hues other than green, or the ability to be photoconverted. To obtain GECIs with these properties, it is generally necessary to undertake the labor-intensive re-engineering of a new GECI, starting from a new parent FP. Notable examples of such efforts include the development of GECIs that are mNeonGreen-derived,^5,6^ yellow fluorescent,^7,8^ red fluorescent,^9–11^ near-infrared fluorescent,^12,13^ or photoconvertible.^14–16^

One GECI feature that has remained under-explored is blue-shifted excitation and emission. Blue-shifted GECIs with anionic chromophore are expected to be much brighter under two-photon excitation^17^ which could enable Ca^2+^ imaging with increased sensitivity. Furthermore, blue-shifted GECIs could be preferred relative to green fluorescent GECIs for applications that combine two-photon activation of opsin-based optogenetic actuators and Ca^2+^ imaging. There is strong overlap of the two-photon spectrum of GCaMPs with the spectra of the most commonly used opsin-based optogenetic actuators, and so there is inevitably unwanted optogenetic activation during Ca^2+^ imaging. In principle, a blue-shifted GECI, with effective two-photon excitation at ∼800 nm, would circumvent this problem. It must be noted that a blue-shifted GECI with performance comparable to a recent generation GCaMP, would still have some inherent disadvantages, such as reduced working depth, due to increased scattering of blue-shifted light when it passes through tissue.

Previous efforts to develop blue-shifted GECIs have relied on the same strategy that was originally used to convert avGFP in a blue FP (BFP) – mutation of the tyrosine residue in the chromophore forming tripeptide to histidine (Y66H).^18^ For example, B-GECO1,^9^ BCaMP1,^19^ and X-CaMP-B,^20^ are blue fluorescent GECIs that were created using this strategy. Unfortunately, these blue fluorescent GECIs suffer from substantially lower sensitivity and lower brightness, relative to optimized GCaMP variants. A blue-shifted Ca^2+^ indicator optimized for Ca^2+^-dependent change in fluorescence lifetime, with a tryptophan-derived chromophore (Y67W), has also been reported.^21^

In contrast to the engineered BFPs with the Y67H mutation, there are naturally occurring blue-shifted FPs that retain a tyrosine-derived chromophore.^22^ One such FP is the tetrameric cFP484 cyan FP (CFP) from *Clavularia sp.*, which was engineered to give the monomeric teal fluorescent protein 1 (mTFP1).^23^ mTFP1 (excitation maximum 462 nm, emission maximum 492 nm) is blue-shifted and 1.6× brighter, relative to avGFP-derived EGFP (excitation maximum 488 nm, emission maximum 508 nm).^1,24^ Molina et al. have demonstrated that blue-shifted FPs with tyrosine-derived chromophores are substantially brighter than EGFP under two-photon excitation.^17^ The promising properties of the mTFP1 parent protein inspired us to attempt to create a new GECI based on this scaffold. Precedent for this effort comes from the successful development of mTFP1-based genetically encoded Zn^2+^ indicators.^25^

In this work, we take advantage of the mTFP1 parent protein to develop a novel GECI named T-GECO1. By capitalizing on the unique spectral properties and high two-photon cross-section of mTFP1, T-GECO1 expands the possibilities for Ca^2+^ imaging experiments and opens new avenues for measuring intracellular Ca^2+^, enabling spectral advantages, compatibility with multiplexing and all-optical experiments, and provides higher two-photon cross-section for enhanced performance *in vitro* and *in vivo*. Here, we present the design, optimization, and characterization of T-GECO1 in soluble protein, cultured neurons, organotypic hippocampal slices, and *in vivo*.

## 2 Methods

### 2.1 Molecular biology and protein engineering

To develop the first prototype of T-GECO1, we fused calmodulin (CaM) and the CaM-binding peptide (CBP) from ncpGCaMP6s to the mTFP1-derived fluorescent protein domain of ZnGreen1.^25,26^ To further improve this prototype, we used multiple rounds of directed evolution. In each round of directed evolution, we initially screened the fluorescence in the context of *Escherichia coli* colonies, selecting for the brightest colonies for further testing. We then cultured these variants and prepared clarified bacterial lysate using B-PER (Thermo Scientific). We measured the fluorescence spectra in the absence of Ca^2+^ (EGTA, buffered in TBS, pH 7.3) and in the presence of 10 nM and 10 mM Ca^2+^ (buffered in TBS, pH 7.3). The DNA encoding variants with improved responses and high brightness was sequenced and used as the template for the next round of library generation.

### 2.2 Protein expression and purification

The pBAD/HisB plasmid carrying the T-GECO1 gene was used to transform chemical or electro-competent *E. coli* DH10B cells which were then grown on solid media. Single colonies were used to inoculate a starter culture supplemented with ampicillin incubated at 37°C. After 4 hours, L-arabinose was added to induce expression, and the culture was shaken overnight at 37°C before harvesting the bacteria by centrifugation. The bacterial pellet was resuspended in 1× TBS, lysed by sonication, and clarified by centrifugation. The cleared lysate was incubated with Ni-NTA resin, washed, and eluted. Dialysis was done into 1× TBS using centrifugal filter units. All steps were carried out at 4°C or on ice, unless specified otherwise.

### 2.3 *In vitro* purified protein characterization

To determine the apparent affinity for Ca^2+^, buffers were prepared with varying concentrations of free-Ca^2+^ ranging from zero to 39 µM by combining appropriate volumes of Ca^2+^-free and Ca^2+^-containing stock solutions.^27^ T-GECO1 was diluted in these buffered solutions, and the fluorescence intensities of the protein in each solution was measured in triplicates. The obtained measurements were plotted on a logarithmic scale against the concentration of free Ca^2+^, and the data was fitted to the Hill equation to determine the apparent *K*d and apparent Hill coefficient.

To measure the extinction coefficient, the Strickler-Berg approach was used.^28^ Briefly, purified T-GECO1 protein was diluted in Ca^2+^-free buffer (30 mM MOPS, 100 mM KCl, 10 mM EGTA, pH 7.2), and Ca^2+^ containing buffer (30 mM MOPS, 100 mM KCl, 10 mM Ca-EGTA, pH 7.2). The absorption, fluorescence emission, and excitation spectra for each sample were collected. For fluorescence measurements, the samples were diluted to have optical densities less than 0.05. Excitation spectra in both samples contain only the contribution from the anionic form of the chromophore. Therefore, we calculated the integral of normalized absorption (entering the Strickler-Berg equation) using corresponding excitation spectra. Fluorescence lifetimes and quantum yields of the anionic chromophore were measured independently and then used in the Strickler-Berg equation.

Fluorescence lifetimes were measured with a Digital Frequency Domain system ChronosDFD appended to a PC1 spectrofluorimeter (both from ISS, Champaign, IL). Fluorescence was excited with a 445-nm laser diode (ISS) through a 440/20 filter. The excitation was modulated with multiple harmonics in the range of 10−300 MHz. Coumarin 6 in ethanol with τ = 2.5 ns (ISS) was used as a lifetime standard to obtain the instrumental response function in each measurement. Fluorescence of the sample and standard were collected at 90° through a 520LP filter to cut off scattered excitation light. The modulation ratio and phase delay curves were fitted to model functions corresponding to a single-or double-exponential fluorescence decay with Vinci 3 software (ISS). Only double exponential decay functions provided acceptable χ^2^ value of 0.5. The main decay component, contributing ∼93% of integrated decay in both samples was used in the Strickler-Berg equation.

Fluorescence quantum yields were determined using the absolute method with an integrating sphere instrument, Quantaurus-QY (Hamamatsu). In this measurement, the quantum yield (QY) was measured as a function of excitation wavelengths between 400 and 500 nm with the step of 5 nm. The quantum yield did not depend on wavelength in the region from 450 – 475 nm for the Ca^2+^-bound state and from 465 – 480 nm for the Ca^2+^-free state, where the anionic absorption dominated. The average of the quantum yields in these regions were calculated and presented in the Results section. All measurements were made in triplicates and averaged.

### 2.4 Two-photon measurements

The two-photon excitation spectra and two-photon absorption cross-sections of T-GECO1 were measured using a previously described protocol.^29^ Briefly, a tunable femtosecond laser (InSight DeepSee, Spectra-Physics, Santa Clara, CA) was coupled to a PC1 Spectrofluorometer (ISS, Champaign, IL). Quadratic power dependence of fluorescence intensity was verified across the spectrum for both proteins and standards. The two-photon cross-section (σ_2_) of the anionic form of the chromophore was determined for both the Ca^2+^-free and Ca^2+^-bound states, as previously described.^30^ As a reference standard, a solution of fluorescein in water at pH 12 was used. Fluorescence intensities of the sample and reference were measured for two-photon excitation at 900 nm and for one-photon excitation at 458 nm (Ar^+^ laser line). Fluorescence measurements utilized a combination of filters (770SP and 520LP). The two-photon absorption spectra were normalized based on the measured σ_2_ values.

### 2.5 Kinetic measurements

Stopped flow kinetic measurements of Ca^2+^ binding and unbinding to T-GECO1 were made using an Applied Photophysics SX20 Stopped-Flow Reaction Analyzer using fluorescence detection. The deadtime of the instrument was 1.1 ms. The mixtures of the protein and Ca^2+^ (or EGTA, for dissociation (or off) rate) were excited at 488 nm with 2 nm bandwidth and the emitted light was collected at 515 nm through a 10-mm path. A total of 10,000 data points were collected over three replicates (n = 3) at increments of 0.01 s for 5 seconds. For the off rate, T-GECO1 (diluted in 5 µM Ca^2+^ in TBS), was rapidly mixed 1:1 with 100 mM EGTA (diluted in TBS). Graphpad Prism 9 was used to fit the decrease in fluorescence intensity observed over time to a single exponential dissociation. The *k*_off_ determined from this fit is the rate constant for dissociation of Ca^2+^ with units of s^-1^. For association (or on) rate, T-GECO1 was diluted in zero free CaEGTA buffer (Thermo Scientific), and mixed 1:1 with varying Ca^2+^ concentrations (150 nM, 225 nM, 351 nM, 602 nM, 1.35 µM). The slope of *k*_obs_ vs. Ca^2+^ concentration was used to determine the *k*_on_ rate (with units of s^-1^M^-1^).

### 2.6 Neuronal stimulation

T-GECO1, GCaMP6s, and jGCaMP8s, were cloned and packaged into AAV2/1 virus under control of the hSyn promoter. The AAVs were used to transduce hippocampal and cortical mixture primary cultures from neonatal (P0) pups in poly-D-lysine-coated 24-well glass bottom plates. After 14 days post transduction, the culture medium was exchanged with 1 mL imaging buffer (145 mM NaCl, 2.5 mM KCl, 10 mM glucose, 10 mM 4-(2-hydroxyethyl)piperazine-1-ethanesulfonic acid (HEPES), 2 mM CaCl_2_, 1 mM MgCl_2_, pH 7.3) containing 10 µM 6-cyano-7-nitroquinoxaline-2,3-dione (CNQX), 10 µM 3-((*R*)-2-carboxypiperazin-4-yl)-propyl-1-phosphonic acid ((*R*)-CPP), 10 µM gabazine, and 1 mM (*S*)-α-methyl-4-carboxyphenylglycine ((*S*)-MCPG) (Tocris). Neurons were field stimulated with 1, 3, 10, and 20 pulses at 30 Hz, and imaged through a 20× objective, with excitation at 470/40 nm. Imaging was performed at room temperature.

### 2.7 Preparation of organotypic hippocampal slice cultures for two-photon Ca^2+^ imaging using T-GECO1 and jGCaMP7s

Organotypic hippocampal slices were prepared from postnatal day 8 (P8) mice (Janvier Labs, C57Bl/6J). Hippocampi were dissected and sectioned into 300 μm thick slices using a tissue Chopper (McIlwain type 10180, Ted Pella), in a cold dissection medium consisting of GBSS (Sigma, G9779) that was supplemented with 25 mM D-glucose, 10 mM HEPES, 1 mM Na-pyruvate, 0.5 mM α-tocopherol, 20 nM ascorbic acid, and 0.4% penicillin/streptomycin (5000 U mL^-1^).

Slices were incubated for 45 minutes at 4 °C in the dissection medium, then placed on a porous membrane (Millipore, Millicell CM PICM03050) and cultured at 37 °C, 5% CO_2_ in a medium consisting of 50% Opti-MEM (Fisher 15392402), 25% heat-inactivated horse serum (Fisher 10368902), 24% HBSS, 1% penicillin/streptomycin (5000 U mL^−1^), and supplemented with 25 mM D-glucose, 10 mM HEPES, 1 mM Na-Pyruvate, 0.5 mM α-tocopherol, 20 nM ascorbic acid, and 0.4% penicillin/streptomycin (5000 U mL^−1^). After three days *in vitro* (DIV), this medium was replaced with one consisting of 82% neurobasal-A (Fisher 11570426), 15% heat-inactivated horse serum (Fisher 10368902), 2% B27 supplement (Fisher, 11530536), 1% penicillin/streptomycin (5000 U mL^−1^), 0.8 mM L-glutamine, 0.8 mM Na-Pyruvate, 10 nM ascorbic acid and 0.5 mM α-tocopherol. This medium was removed and replaced every 2-3 days. Slices were transduced with AAVs at DIV 3 by bulk application of 1 µL of virus per slice, for expression of T-GECO1 or jGCaMP7s under control of the hSyn promoter. Experiments were performed at DIV 10.

### 2.8 Two-photon Ca^2+^ imaging of action potentials in T-GECO1-and jGCaMP7s-expressing organotypic hippocampal slices

At DIV 10, whole-cell current clamp recordings of T-GECO1- or jGCaMP7s-expressing neurons were performed at room temperature (21 – 23°C). A commercial upright microscope (Zeiss, Axio Examiner.Z1), equipped with a microscope objective (Zeiss, W Plan-Apochromat 20X, 1.0 NA) and an sCMOS camera (Photometrics, Kinetix), was used to collect light transmitted through the sample. Patch-clamp recordings were performed using an amplifier (Molecular Devices, Multiclamp 700B) and a digitizer (Molecular Devices, Digidata 1440A), at a sampling rate of 10 kHz using pCLAMP10 (Molecular Devices). During the experiments, slices were continuously perfused with artificial cerebrospinal fluid (ACSF) composed of 125 mM NaCl, 2.5 mM KCl, 1.5 mM CaCl_2_, 1 mM MgCl_2_, 26 mM NaHCO_3_, 0.3 mM ascorbic acid, 25 mM D-glucose, 1.25 mM NaH_2_PO_4_. ACSF was supplemented with 1 μM AP5 (Abcam, ab120003), 1 μM NBQX (Abcam, ab120046), and 10 µM picrotoxin (Abcam, 120315). Continuous aeration of the recording solution with 95% O_2_ and 5% CO_2_ resulted in a pH of 7.4. Patch pipettes were pulled from borosilicate glass capillaries (with filament, OD: 1.5 mm, ID: 0.86 mm, 10 cm length, fire polished, WPI) using a Sutter Instruments P1000 puller, to a tip resistance of 4.5 – 5.5 MΩ, and filled with an intracellular solution consisting of 135 mM K-gluconate, 4 mM KCl, 4 mM Mg-ATP, 0.3 mM Na_2_-GTP, 10 mM Na_2_-phosphocreatine, and 10 mM HEPES (pH 7.35). Only recordings with an access resistance below 20 MΩ were included in subsequent analysis. In the current-clamp configuration, the bridge potential was corrected (bridge potential = 13.9 ± 1.0 MΩ; mean ± s.d.). Two-photon scanning imaging was performed with a Ti:sapphire tunable pulsed laser (Spectra Physics, Mai-Tai DeepSee, pulse width ≈ 100 fs, repetition rate 80 MHz, tuning range 690 – 1040 nm), going through a commercial galvo-galvo scanning head (3i, Vivo 2-photon) operated using Slidebook 6 software. The detection axis consisted of a PMT with a 510/84 nm bandpass filter (Semrock, FF01-510/84). Imaging was performed within a 365 × 365 µm field of view (FOV) at a rate of 3.05 Hz (bidirectional scanning, 256 × 256 pixels, pixel size 1.4 µm, dwell time 4.0 µs). Laser power was controlled by a Pockels cell (Conoptics, 350-80). Prior to the experiments, powers were measured at the output of the objective using a thermal sensor power meter (Thorlabs, PM100D).

Action potentials were triggered by injecting current for 5 ms (ranging from 500 to 1200 pA), at a rate of 30 Hz during a period ranging from 5 ms to 650 ms, in order to evoke the desired number of action potentials, while the FOV was scanned under 850 nm or 920 nm illumination at 20 mW. Recordings were dismissed if the desired amount of action potentials failed to occur.

Fluorescence intensities were integrated over regions of interest (ROI) covering the patched neuron soma. Percentage changes in fluorescence were calculated as /¢i*F*/*F*_0_ = (*F* – *F*_0_)/*F*_0_, where *F*_0_ is the basal level of fluorescence measured, averaged over 35 frames (≈ 12 s) before the triggering of action potentials. SNR was measured as *SNR* = *F*/σF_0_, where σF_0_ represents the standard deviation of the fluorescence *F* over the 35 frames prior to the stimulation.

### 2.9 Evaluation of crosstalk induced by the two-photon scanning laser in ChroME-expressing organotypic hippocampal slices

At DIV 3, organotypic hippocampal slices were infected with a mixture of AAV9.hSyn.DIO.ChroME.Flag.ST.P2A.H2B.mRuby3.WPRE.SV40 (titer = 5.9E12 GC mL^-1^) and AAV9.hSyn.Cre.WPRE.hGH (titer = 2.3E11 GC mL^-1^) by bulk application of 1 µL of the mixture.

At DIV 10, whole-cell current clamp recordings of ChroME-expressing cells were performed in the same conditions as described above. The membrane potential of the patched neuron was monitored and recorded while scanning the FOV for 30 s (365 × 365 µm^2^, 256 × 256 pixels, pixel size 1.4 µm) at 850 nm or 920 nm, at 20 mW, and at acquisition rates of 1.5 Hz, 3.05 Hz and 6 Hz (corresponding to dwell time per pixel of 6 µs, 4 µs and 2 µs respectively). The variation of membrane potential Δ*V*_m_ reported in the manuscript corresponds to the average of the amplitude of the depolarization peaks induced by the imaging laser, during a 30 s scanning epoch. Depolarization peaks were measured as/Δ*V*_m_ = *V*_mp_ – *V*_m0_, where *V*_mp_ is the peak of the membrane potential depolarization (one for each frame) and *V*_m0_ is the membrane potential of the neuron measured just before the beginning of the scanning. The ratio Δ*V*_m850_/Δ*V*_m920_, was calculated for each cell, and then averaged across cells.

### 2.10 Stereotaxic injection and fiber implant surgery

Stereotaxic injections of AAVs and optical fiber implant surgeries were performed at the same time in C57BL/6J mice (The Jackson Laboratory, #000664) at around P60. Mice were anesthetized with isoflurane and monitored throughout the surgery with tail pinch and breathing rate. First, the skin above the skull was cleaned and removed to allow attachment of the headframe and optical fiber implants. Next, a burr hole craniotomy was drilled above the fiber implant coordinates for implantation in the nucleus accumbens core (AP: 1.2 mm, ML: 1.3 mm, DV: 4.1 mm). Virus injection of either AAV2/1-hSyn-T-GECO1 (100 nL, titer = 1.5E13 GC mL^-1^) or AAV2/1-hSyn-jGCaMP8s (100 nL, titer = 1.9E13 GC mL^-1^) was performed with a glass pipette prior to fiber implant. Following virus injection, a fiber optic probe was positioned above the same coordinates and the tip of the fiber was lowered to 100 μm above the virus injection. The fiber implant was then affixed to the skull with dental cement. A custom headframe was then positioned on the skull and glued in place with dental cement to allow head-fixation during photometry. The mice were allowed to recover for two weeks before the start of imaging. All photometry was performed in head-fixed mice placed on a running wheel to allow spontaneous running.

### 2.11 Fiber photometry measurement and analysis

Fiber photometry measurements were performed on a custom spectral photometry system. 448 nm (Coherent, OBIS 445 nm LX 365 mW LASER, measured wavelength is 448 nm) and 473 nm (Coherent, OBIS 473 nm LX 200 mW LASER, measured wavelength is 473 nm) excitation lasers were co-aligned and focused onto the back pupil of an objective (Nikon, Plan Apochromat, 10X, 0.45 NA, 25 mm FOV). The excitation light was coupled into a fiber optic patch cable (Doric, 200 μm core, 0.37 NA) by positioning the patch cable at the image plane of the objective. The other end of the patch cable was coupled to the implanted fiber stub. Emitted light from the brain tissue was collected through the same fiber probes and patch cable and passed through a polychromator (Edmund Optics, 50 mm N-SF11 equilateral prism). The polychromator spreads the image of the fiber tip according to its spectrum, which was imaged onto a sCMOS camera sensor (Hamamatsu, Orca Flash 4.0 v3). The excitation lasers and camera sensor were triggered synchronously using an Arduino Teensy board (the excitation lasers were sequentially triggered while the camera sensor was triggered at every frame) at 24 Hz frame rate. The raw images were acquired and saved through a custom script in the Bonsai reactive programming environment.^31^ The recorded spectra corresponding to either T-GECO1 (485-510 nm) or jGCaMP8s (520-545 nm) emission were averaged to yield a single intensity time trace. The fractional intensity change was computed by dividing the intensity of each frame by the mean fluorescence of the full trace over time.

### 2.12 Animal care

Animal experiments at Sorbonne Université were conducted in accordance with guidelines from the European Union and institutional guidelines on the care and use of laboratory animals (Council Directive 2010/63/EU of the European Union). Surgery protocol and fiber photometry imaging experiments at the Allen Institute for Neural Dynamics were approved by the Allen Institute Institutional Animal Care and Use Committee (IACUC). Animal experiments at Janelia Research Campus were conducted according to National Institutes of Health guidelines for animal research and were approved by the Janelia Research Campus Institutional Animal Care and Use Committee and Institutional Biosafety Committee. Procedures in the United States conform to the National Institutes of Health (NIH) Guide for the Care and Use of Laboratory Animals. Mice were housed under controlled temperature (approximately 21 °C) and humidity (approximately 50%) conditions under a reverse light cycle.

## 3 Results

### 3.1 Development of mTFP1-based genetically encoded Ca^2+^ indicator, T-GECO1

Our initial template for constructing an mTFP1-based GECI was the mTFP1-based genetically encoded Zn^2+^ indicator, ZnGreen1.^25^ ZnGreen1 consists of the Zap1 zinc finger inserted into a further engineered version of mTFP1. This version of mTFP1 in ZnGreen1 harbors the nine additional mutations N42H, N81D, D116G, S146C, T147D, R149K, E168K, R198H, V218A using mTFP1 numbering (or N42H, N81D, D116G, S323C, T324D, R326K, E345K, R375H, V395A using T-GECO numbering). To construct the initial prototype mTFP1-based GECI, designated T-GECO0.1, we replaced the Zap1 zinc finger of ZnGreen1 with the fused calmodulin (CaM) and CaM-binding peptide (CBP) domain from ncpGCaMP6s.^26^ The linker sequences from ZnGreen1 were retained. The arrangement of these domains is represented in **Fig. 1a**.

As previously described, we define the linkers as additional residues that are inserted between the Ca^2+^-binding domain (CaM fused to CBP) and the gatepost residues 143 and 146 of mTFP1.^32^ In T-GECO0.1 the linker from the first mTFP1 gatepost (W143) to CaM linker (Linker 1) is Leu-Gly-Asn. Linker 2 from CBP to the second mTFP1 gatepost (S146C) is a single Pro. To develop further improved T-GECO variants, we first optimized these linker residues and some adjacent positions. This was achieved by randomizing each residue, expressing the resulting library in *E. coli*, picking and culturing bright colonies, and testing Ca^2+^-dependent responses in bacterial lysates. Ultimately, we identified Arg-Asn-Arg as the optimal Linker 1, and Ile as the optimal Linker 2 (**Fig. 1a**).

Further optimization by directed evolution was performed by generating libraries using error-prone (EP) PCR amplification of the entire coding sequence of T-GECO. In each round, we took variants with moderate to high fluorescence change upon binding Ca^2+^ and measured their affinity, pH response, quantum yield, and extinction coefficient. We obtained the DNA sequence of these variants and used them as the template for the next round of iterative directed evolution. Following five generations of screening, we arrived at T-GECO1 on the basis of its high Δ*F*/*F*_0_, high affinity, high brightness, two-photon cross-section, and kinetics. T-GECO1 has 25 mutations with respect to T-GECO0.1 (E5D, T96I, H123Y, L144R, G145N, N146R, K175E, E199V, T207A, A218S, K222N, R251H, H252R, T262S, E268D, M269V, Q280L, R304H, P322I, C323G, D324G, K339E, K341E, T400R, D401R, using T-GECO numbering). There are 4 mutations in the linkers, 9 mutations in the mTFP1-derived region, 11 mutations in CaM, and 1 mutation in CBP. The locations of all mutations are shown in **Fig. 1a** and **Fig. 2**.

We first characterized the photophysical properties of T-GECO1 as a soluble protein under one-photon and two-photon excitation (**Fig. 1b-f**). Under one-photon excitation, T-GECO1 in the Ca^2+^-bound state exhibits excitation and emission peaks at 468 nm and 500 nm, respectively. The molecular brightness of T-GECO1 in the Ca^2+^-bound state, calculated as the product of the extinction coefficient (49,300 M^-1^cm^-1^) and quantum yield (0.83), is similar to that of EGFP (**Table 1**).^17^ The two-photon excitation maximum of T-GECO1 is 888 nm with a brightness of 83 GM, which is 1.4× the value of mTFP1 and 2× the value of EGFP (**Table 2**). T-GECO1 exhibits a large change in fluorescence intensity upon addition of Ca^2+^, with 1-photon peak Δ*F*/*F*_0_ of 15 and 2-photon peak Δ*F*/*F*_0_ of 7, where Δ*F*/*F*_0_ = (*F*_max_ – *F*_min_)/*F*_min_. Additionally, we determined that T-GECO1 has an apparent *K*_d_ of 82 nM for binding to Ca^2+^, and an apparent Hill coefficient (n_H_) of

### 3.6. T-GECO1 exhibits moderate binding (on) and dissociation (off) kinetics as soluble protein, with *k*_on_ of 8.5 × 10^5^ M^-1^s^-1^ and *k*_off_ of 1.02 s^-1^

Together, these results demonstrate T-GECO1 has favorable photophysical characteristics that make it a potentially useful new GECI. Its high fluorescence change upon binding Ca^2+^, high brightness and two-photon cross-section, and reasonable association and dissociation kinetics suggest that T-GECO1 is a promising tool for monitoring Ca^2+^ dynamics using blue-shifted excitation.

**Fig. 1.**
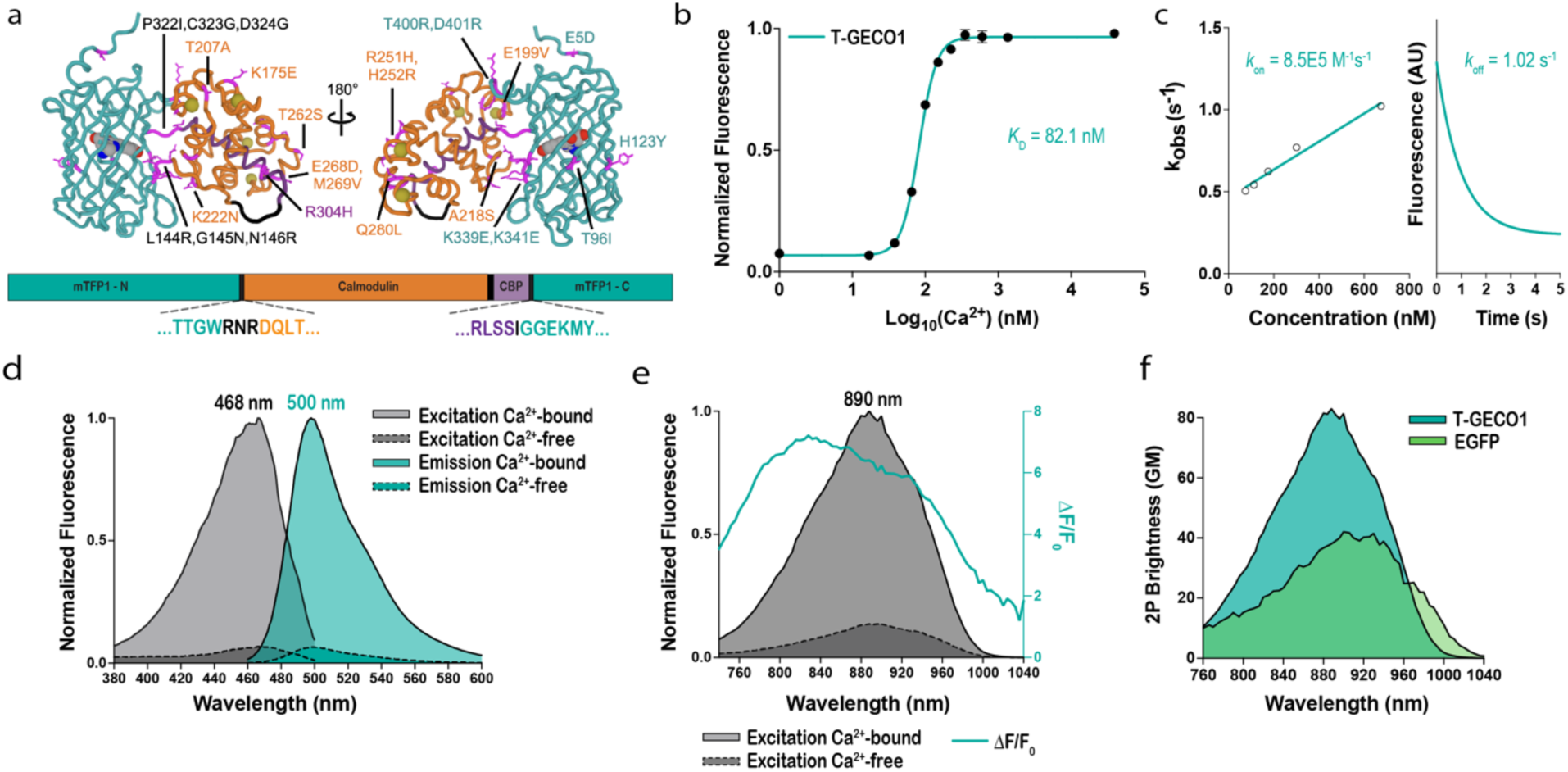
Development and characterization of T-GECO1 as a purified protein. (**a**) Two views of the modeled structure of the Ca^2+^-bound state of T-GECO1. For the structure representation, mutated residues are shown as magenta sticks, Ca^2+^ is shown as yellow spheres, and the chromophore is shown in space-filling representation. Both the protein structure and the labels are shown in teal for the mTFP1-derived domain, in orange for the CaM domain, in purple for the CaM-binding peptide, and in black for linkers. Colors are consistent with the sequence alignment shown as Fig. 2. The overall structure was predicted using ColabFold.^33^ The chromophore was positioned using PyMol (Version 2.5.4 Schrödinger, LLC.) to superimpose the structure of mTFP1 (PDB ID 2HQK)^23^ with the fluorescent protein portion of the T-GECO1 model. Ca^2+^ ions were similarly positioned by superimposing the CaM domain of GCaMP2 (PDB ID 3EVR)^34^ with the CaM portion of the T-GECO1 model. (**b**) Ca^2+^ titration of T-GECO1. (**c**) Stopped-flow kinetic measurements of the fluorescence response of T-GECO1 for Ca^2+^ association (left) and dissociation (right). (**d**) Excitation and emission spectra of T-GECO1 in the presence and absence of Ca^2+^. (**e**) Two-photon excitation-induced fluorescence of T-GECO1 as a function of wavelength, in the presence and absence of Ca^2+^, with Δ*F*/*F*_0_ represented in teal. (**f**) Two-photon cross-section of T-GECO1 in the Ca^2+^-bound state, compared to the two-photon cross-section of EGFP.

**Fig. 2.**
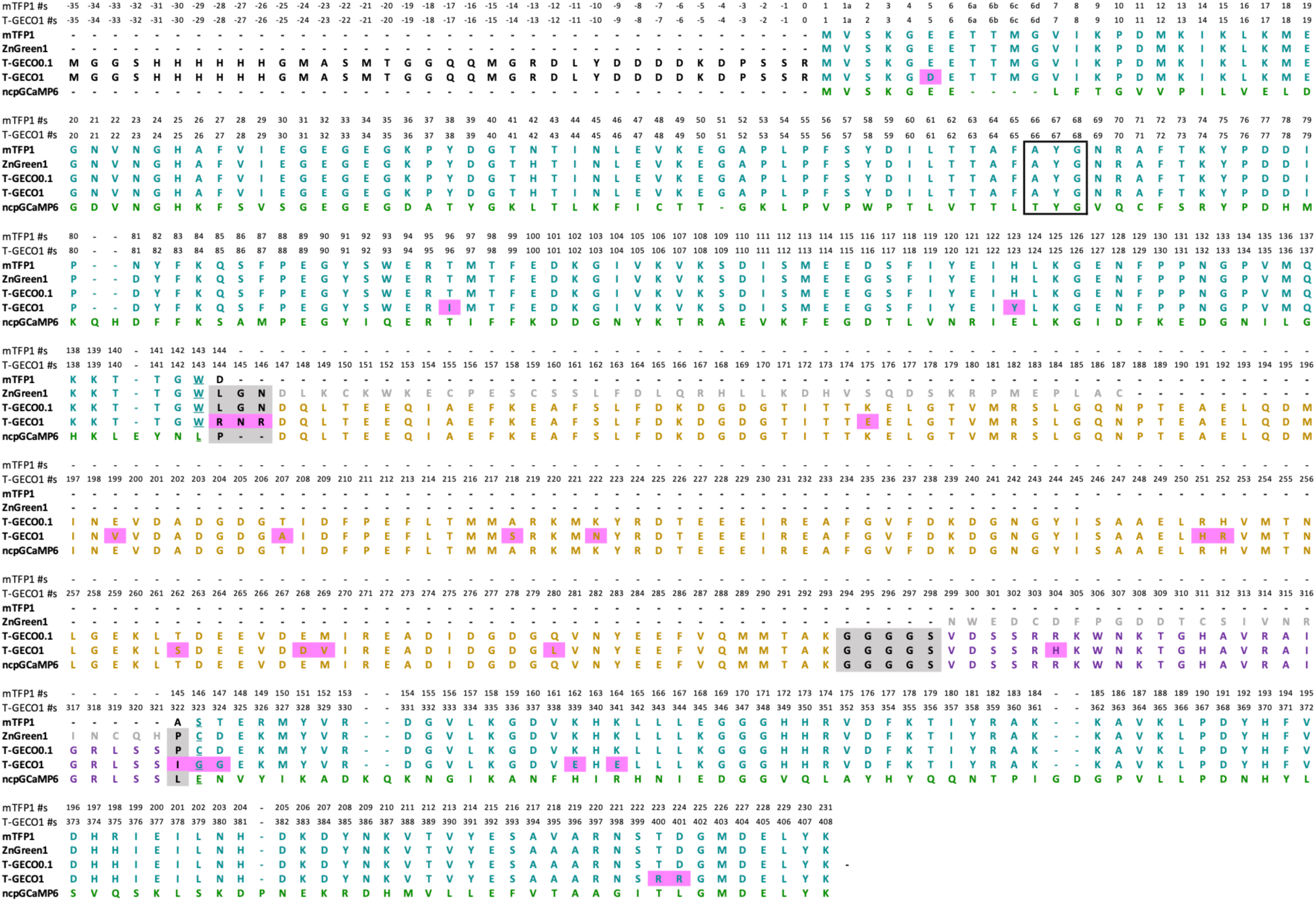
Sequence alignment of T-GECO1 and related proteins. Residues are colored teal for the mTFP1-derived domain, orange for the CaM domain, purple for the CaM-binding peptide, and black on a gray background for the linkers. Mutated residues are shown on a magenta background. A black box encloses the chromophore-forming tripeptide. The gatepost residues 143 and 146 (using mTFP1 numbering) are underlined.^32^ Colors are consistent with the structural model shown in Fig. 1a.

**Table 1.**
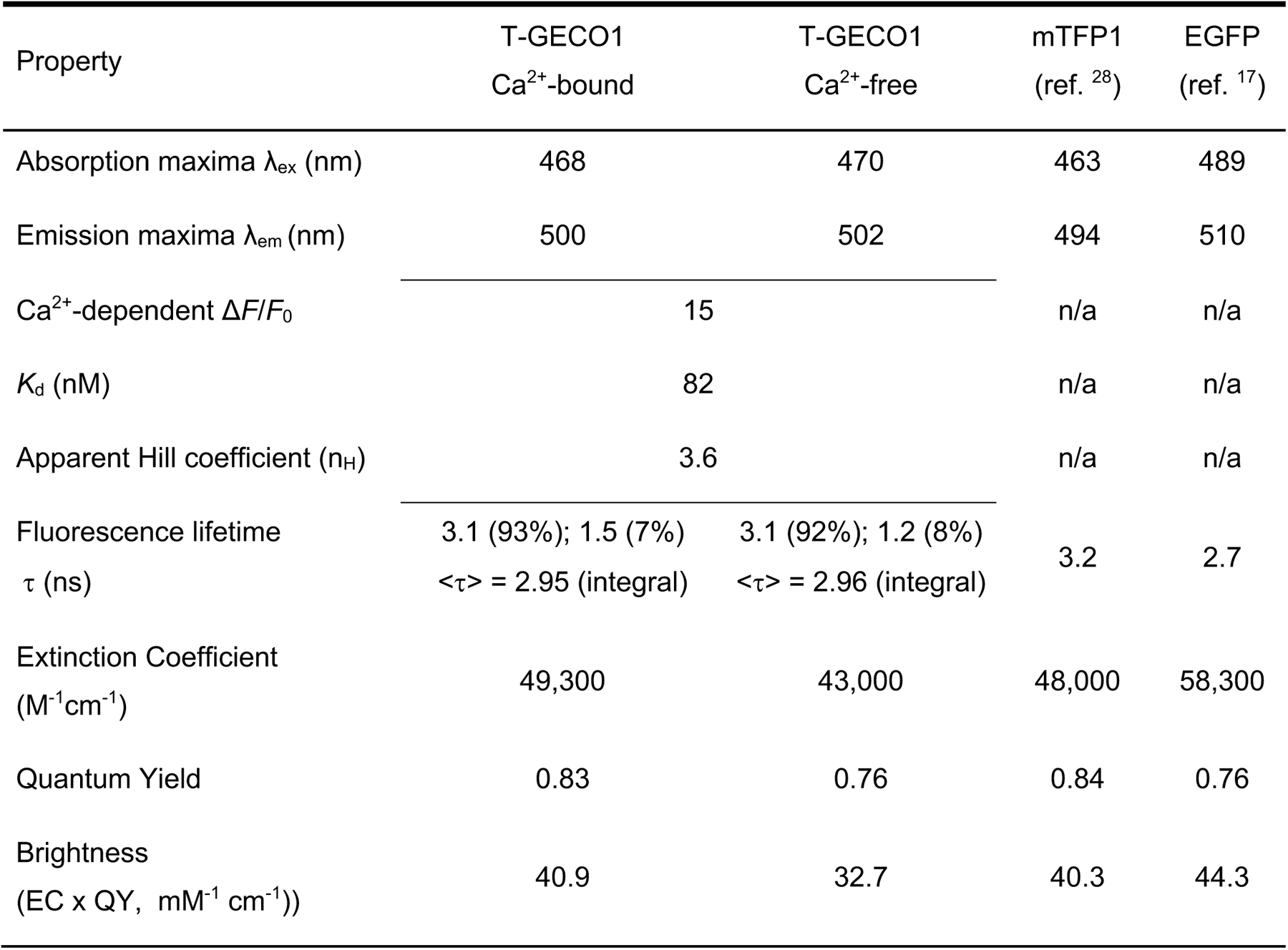
One-photon photophysical properties of T-GECO1 and mTFP1,^28^ measured as purified proteins (n = 3, averaged). Extinction coefficients were obtained using Strickler-Berg formula.^28^ In this calculation, the main fluorescence lifetime component of T-GECO1 was used. Note that the relative values of the brightness of the Ca^2+^-bound and Ca^2+^-free fluorescent states shown here do not represent the Ca^2+^-dependent fluorescence change of T-GECO1. The Ca^2+^-dependent fluorescence change is primarily due to a change in the protonation state of the chromophore which changes the fraction of the protein in fluorescent state.

| Property | T-GECO1<br>Ca<sup>2+</sup>-bound | T-GECO1<br>Ca<sup>2+</sup>-free | mTFP1<br>(ref. <sup>28</sup>) | EGFP<br>(ref. <sup>17</sup>) |
| --- | --- | --- | --- | --- |
| Absorption maxima $\lambda_{ex}$ (nm) | 468 | 470 | 463 | 489 |
| Emission maxima $\lambda_{em}$ (nm) | 500 | 502 | 494 | 510 |
| Ca<sup>2+</sup>-dependent $\Delta F/F_0$ | 15 | | n/a | n/a |
| $K_d$ (nM) | 82 | | n/a | n/a |
| Apparent Hill coefficient ($n_H$) | 3.6 | | n/a | n/a |
| Fluorescence lifetime<br>$\tau$ (ns) | 3.1 (93%); 1.5 (7%)<br><$\tau$> = 2.95 (integral) | 3.1 (92%); 1.2 (8%)<br><$\tau$> = 2.96 (integral) | 3.2 | 2.7 |
| Extinction Coefficient<br>(M<sup>-1</sup>cm<sup>-1</sup>) | 49,300 | 43,000 | 48,000 | 58,300 |
| Quantum Yield | 0.83 | 0.76 | 0.84 | 0.76 |
| Brightness<br>(EC x QY, mM<sup>-1</sup> cm<sup>-1</sup>) | 40.9 | 32.7 | 40.3 | 44.3 |

**Table 2.** Two-photon photophysical properties of T-GECO1 (Ca^2+^-bound state), mTFP1, and EGFP, measured as purified proteins (n = 3, averaged). Values for mTFP1 and EGFP have were previously reported.^17,28^

| Property | T-GECO1<br>Ca<sup>2+</sup>-bound | T-GECO1<br>Ca<sup>2+</sup>-free | mTFP1 | EGFP |
| --- | --- | --- | --- | --- |
| Two-photon cross-sections<br>(GM) with $\lambda_{\text{max}}$ in parentheses | 100 (888 nm) | 82 (896) | 70 (875 nm) | 54 (911 nm) |
| Two-photon brightness $F_2$<br>(GM) with $\lambda_{\text{max}}$ in parentheses | 83 (888 nm) | 62 (896) | 60 (875 nm) | 41 (911 nm) |
| Two-photon $\Delta F/F_0$ | 7.4 | | n/a | n/a |

### 3.2 Imaging of Ca^2+^ in electric field stimulated neuronal cultures

To characterize T-GECO1 in neuronal cultures using one-photon excitation (excitation at 450 – 490 nm), we expressed it under the control of human synaptic (hSyn) promoter in rat primary cortical and hippocampal neurons. We compared the performance of T-GECO1 to GCaMP6s and jGCaMP8s (**Fig. 3a**). To evoke neuronal activity, we applied trains of 1, 3, 10, and 20 electric field stimuli and analyzed the resulting fluorescence changes (**Fig. 3b,c,d**). T-GECO1 exhibited a peak change in fluorescence (Δ*F*/*F*_0_) of 3% for a single stimulus. In comparison, GCaMP6s and jGCaMP8s had peak responses of 9% and 20%, respectively, in response to single stimuli. T-GECO1 exhibited lower Δ*F*/*F*_0_ values across all numbers of stimuli tested. The baseline brightness of T-GECO1 (536 +/-59 RFU), was found to be 16% higher than that of GCaMP6s (463 +/-16 RFU) and 15% higher than that of jGCaMP8s (466 +/-21 RFU) (**Fig. 3e**). T-GECO1 exhibited a marginally larger SNR compared to GCaMP6s and jGCaMP8s, partially due to its higher baseline brightness (**Fig. 3f**).

These results demonstrate that T-GECO1 has sufficient sensitivity for detecting small numbers of action potentials in cultures, using one-photon excitation. However, further optimization will be necessary to achieve the peak sensitivity exhibited by late-generation GCaMP series indicators. Nevertheless, T-GECO1’s higher baseline brightness and blue-shifted excitation and emission may prove advantageous, relative to the GCaMP series, for certain one-photon excitation applications such as multicolor imaging and combined use with longer-wavelength activatable optogenetic tools.

**Fig. 3.**
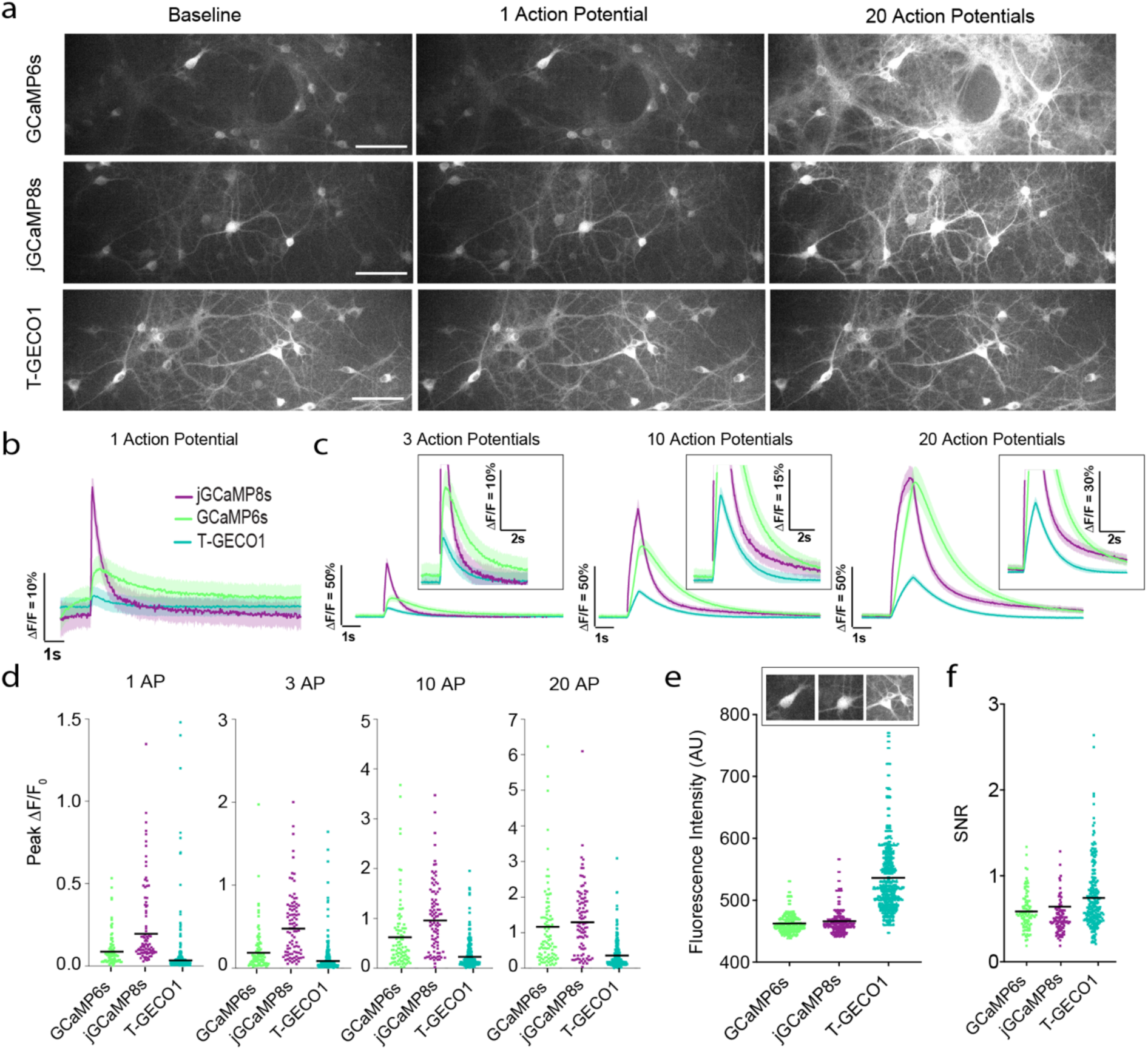
Characterization of T-GECO1 in rat cultured neurons. (**a**) Images of primary rat hippocampal cultured neurons expressing GCaMP6s, jGCaMP8s, and T-GECO1 under the hSyn promoter at baseline, and after field stimulations of 1 and 20 action potentials (APs) at room temperature. (**b**) Normalized Δ*F*/*F*_0_ traces for stimulations at 1 AP and (**c**) 3 AP, 10 AP, and 20 AP at 30 Hz. (**d**) Peak Δ*F*/*F*_0_ of the three sensors across the same conditions. (**e**) Baseline brightness of the three sensors. (**f**) Signal-to-noise ratio (SNR) for the three variants across conditions. Traces and error bars denote mean +/-s.e.m. Each data point is one ROI and is pooled across three independent wells.

### 3.3 Two-photon Ca^2+^ imaging of T-GECO1 in organotypic hippocampal slices

Next, we compared the performance of T-GECO1 to jGCaMP7s (ref. ^3^) using two-photon Ca^2+^ imaging in neonatal mouse organotypic hippocampal slices (**Fig. 4a**). We hypothesized that, due to its blue-shifted two-photon excitation maxima relative to GCaMP7s’s (**Fig. 1f**), T-GECO1 could be the more suitable choice for all-optical stimulation and imaging when used in conjunction with the ChroME opsin.^35^ Specifically, we expected that excitation wavelengths that are near-optimal for T-GECO1 (i.e., ∼850 nm) would result in less undesirable activation of ChroME than excitation wavelengths that are near-optimal for GCaMP7s (i.e., ∼920 nm). To test this hypothesis, we quantified the change in membrane potential when ChroME-expressing neurons were illuminated with either 850 nm or 920 nm and expressed the ratio calculated as ΔV_m850_/ΔV_m920_, on a cell-by-cell basis. Across all tested frequencies (1.5, 3, and 6 Hz), this ratio consistently remained below one (0.62, 0.73, 0.67), indicating that using an imaging wavelength of 850 nm rather than 920 nm is advantageous for reducing undesirable ChroME activation (**Fig. 4b**).

We next investigated the fluorescence responses of both T-GECO1 and GCaMP7s at excitation wavelengths of 850 nm and 920 nm, with varying numbers of stimulated action potentials (APs) (**Fig. 4a**). Under excitation at 850 nm, T-GECO1 exhibited a fluorescence change (Δ*F*/*F*_0_) of 73%, whereas jGCaMP7s exhibited a change of 13%, in response to 1 AP (**Fig. 4c,e**). In response to 20 APs, T-GECO1 exhibited a fluorescence change of 450% and jGCaMP7s exhibited a change of 38% (**Fig. 4c,e**). The signal-to-noise ratio (SNR) for T-GECO1 was substantially higher than for jGCaMP7s (**Fig. 4f**).

When excited at 920 nm, the differences between the two indicators were marginal. At 1 AP, both T-GECO1 and jGCaMP7s displayed similar Δ*F*/*F*_0_ values (65% and 75%, respectively). For 20 APs, T-GECO1 exhibited a Δ*F*/*F*_0_ of 547%, approximately 2.8 times greater than the Δ*F*/*F*_0_ of jGCaMP7s (194%) (**Fig. 4g, i**). The baseline brightness of T-GECO1 before stimulation was higher than that of jGCaMP7s (53.6 AU for T-GECO1 and 37.5 AU for jGCaMP7s) and remained higher at its peak after stimulation (88.1 AU and 79.9 AU, respectively) (**Fig. 4h**). Similar to the 850 nm excitation, the SNR of T-GECO1 was consistently higher than that of jGCaMP7s (**Fig. 4j**).

These results indicate that T-GECO1 may offer substantial performance advantages relative to jGCaMP7s under two-photon excitation conditions, particularly at the excitation wavelength of 850 nm. This apparent advantage is consistent with our original rationale for using mTFP1, which is itself particularly bright under two-photon excitation, as the starting point for developing a new Ca^2+^ indicator.

**Fig. 4.**
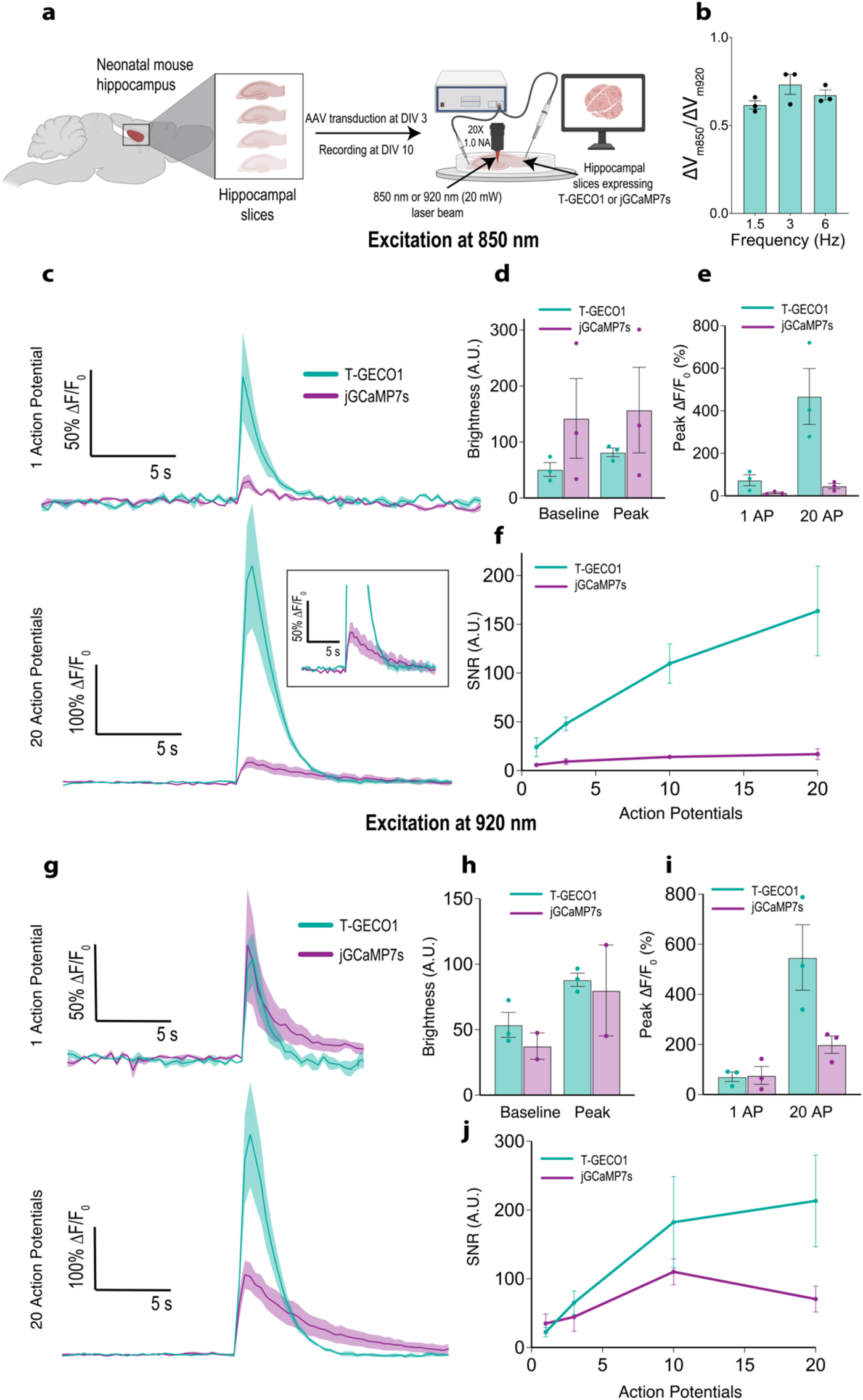
Two-photon Ca^2+^ imaging of T-GECO1 in organotypic hippocampal slices. (**a**) Schematic of the setup. (**b**) ΔV_m850_/ΔV_m920_ ratio of ChroME-expressing organotypic hippocampal slices. (**c**) Representative traces of T-GECO1 (teal) and jGCaMP7s (purple) for 1 action potential (top) and 20 action potentials (bottom) excited at 850 nm. (**d**) Baseline brightness (A.U) for the two indicators at baseline (before stimulation) and at peak (maximum brightness after stimulation) at 850 nm excitation. (**e**) Peak Δ*F*/*F*_0_ (%) for the two indicators at 1 or 20 action potentials at 850 nm excitation. (**f**) SNR (signal-to-noise ratio) for the two indicators with respect to action potentials at 850 nm excitation. (**g**) Representative traces of T-GECO1 (teal) and jGCaMP7s (purple) for 1 action potential (top) and 20 action potentials (bottom) excited at 920 nm. (**h**) Baseline brightness (A.U) for the two indicators at baseline (before stimulation) and at peak (maximum brightness after stimulation) at 920 nm excitation. (**i**) Peak Δ*F*/*F*_0_ (%) for the two indicators at 1 or 20 action potentials at 920 nm excitation. (**j**) SNR (signal-to-noise ratio) for the two indicators with respect to action potentials at 920 nm excitation. Error bars denote +/-S.E.M.

### 3.4 *In vivo* Ca^2+^ detection in the nucleus accumbens using fiber photometry

To evaluate the performance of T-GECO1 in the intact brain using one-photon excitation, we conducted fiber photometry measurements by expressing either T-GECO1 or jGCaMP8s in the nucleus accumbens of mice. Fluorescence traces were recorded using fiber implants positioned above the injection site (**Fig. 5a,b**). We excited both T-GECO1 and jGCaMP8s using either 448 nm or 473 nm wavelengths while the mice engaged in spontaneous running, with occasional manual whisker flicking to evoke Ca^2+^ transients. T-GECO1 enabled reliable detection of Ca^2+^ transients at both 473 nm and 448 nm excitation wavelengths, with higher fluorescence changes (Δ*F*/*F*_0_) observed at 473 nm compared to 448 nm (**Fig. 5c,d**). In general, these responses were substantially lower than those observed with jGCaMP8s, regardless of the excitation wavelength used (**Fig. 5e,f**).

These *in vivo* imaging results are qualitatively consistent with the results from *in vitro* imaging in neuronal cultures, using one-photon excitation. That is, T-GECO1 can be effectively utilized for *in vivo* one-photon excitation imaging of neuronal activity using either 448 nm or 473 nm excitation, but does not achieve the peak sensitivity exhibited by late-generation GCaMP series indicators.

**Fig. 5.**
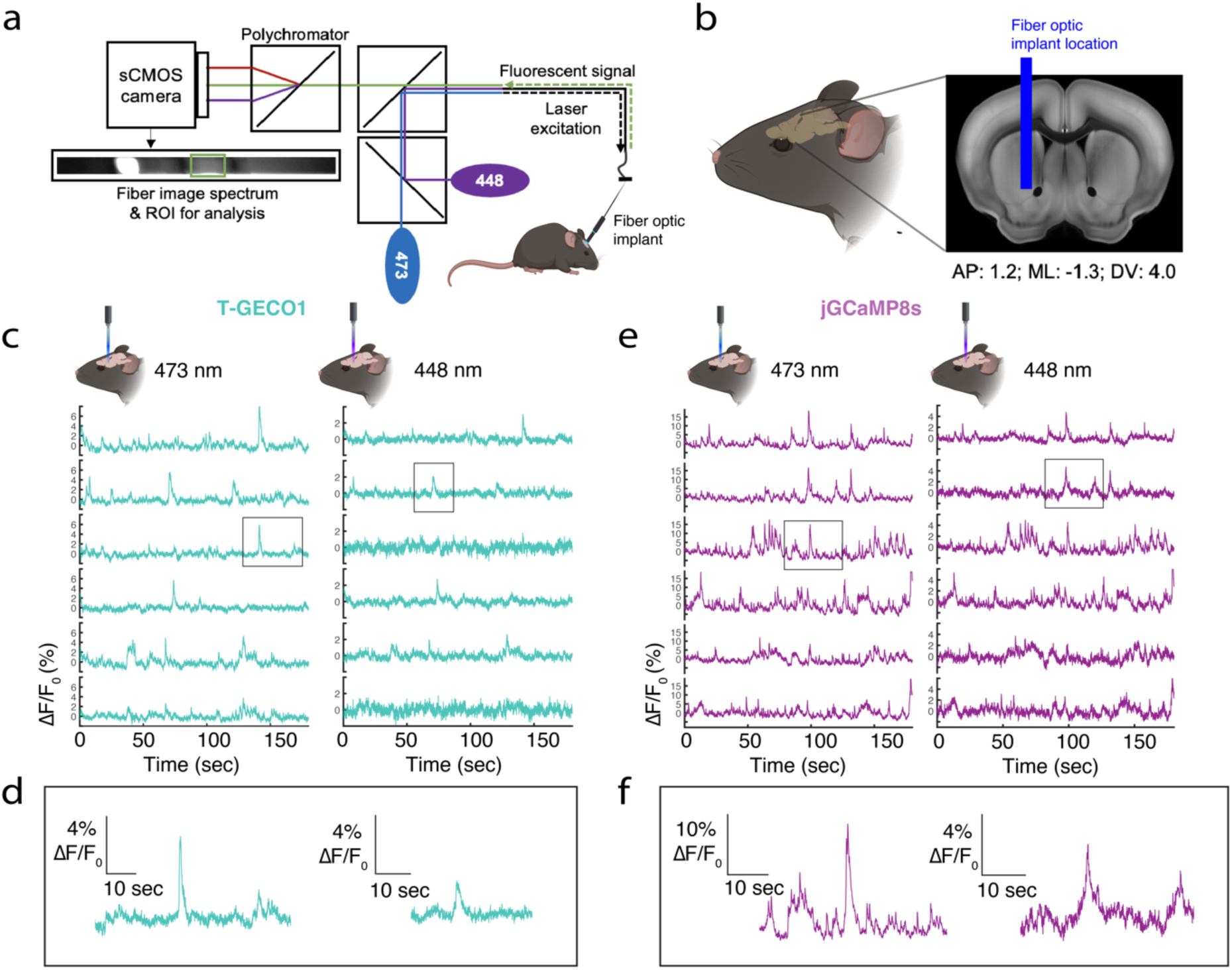
*In vivo* Ca^2+^ detection in the nucleus accumbens using fiber photometry. (**a**) Simplified schematic illustrating the fiber photometry setup, featuring two excitation wavelengths of 473 nm and 448 nm. (**b**) Precise position of the fiber optic implant. (**c**) Representative fluorescence traces of T-GECO1 at 473 nm excitation (left) and 448 nm excitation (right). (**d**) Zoomed-in view of the outlined traces displayed in (**c**). (**e**) Representative fluorescence traces of jGCaMP8s at 473 nm excitation (left) and 448 nm excitation (right). (**f**) Zoomed-in view of the outlined traces displayed in (**e**).

## 4 Discussion

To expand the GECI color palette we developed a novel Ca^2+^ indicator, T-GECO1, based on mTFP1. In this manuscript, we have reported the development and characterization of T-GECO1, and compared it against state-of-the-art GCaMP series indicators for imaging of neuronal activity. We performed this comparison in three different contexts: *in vitro* one-photon excitation in cultured rat hippocampal neurons, *in vivo* one-photon excitation fiber photometry in mice, and *ex vivo* two-photon Ca^2+^ imaging in hippocampal slices.

As we had hoped when we embarked on the development of this new Ca^2+^ indicator, T-GECO1 retains the blue-shifted spectral profile of mTFP1 and its high two-photon cross-section. The results from two-photon imaging in hippocampal slices reveal that these properties can provide a substantial SNR improvement relative to late generation GCaMP variants, particularly for two-photon excitation at 850 nm. We have demonstrated that these properties allow for the reduction of cross-talk in all-optical experiments by reducing the unintended activation of opsins. Other applications that could benefit from the excitability of T-GECO1 at 850 nm could include its combination with red-shifted GECIs or GEVIs to monitor responses from two distinct neuronal populations. Under one-photon excitation conditions, T-GECO1 proved to an effective indicator when measured either in cultured neurons or *in vivo* using fiber photometry.

Based on the precedent of the GCaMP series, further engineering and optimization of T-GECO1 is likely to provide improved versions that will continue to surpass the GCaMP series under two-photon excitation conditions and may one day rival or surpass the GCaMP series under one-photon excitation conditions.

## 5 Conclusion

T-GECO1 is a high-performance first-generation GECI that is an effective blue-shifted alternative to green and red-emitting indicators like jGCaMP8 or the R-GECO1-derived jRGECO1a, respectively.^4,10^ While further rounds of directed evolution and optimization may be necessary to reach the peak sensitivity and responses of the highly optimized GCaMP series under one-photon excitation, the combination of its teal coloration and high two-photon cross-section make T-GECO1 a practically useful new tool for imaging of dynamic changes in Ca^2+^ concentration using two-photon excitation.

## Material and Data Availability

The data supporting this research are available upon request by contacting REC. Plasmid constructs encoding T-GECO1 are available through Addgene or by contacting REC.

## Acknowledgments

This work was supported by grants from the Canadian Institutes of Health Research (CIHR, FS-154310 to REC) and the Natural Sciences and Engineering Research Council of Canada (NSERC, RGPIN 2018-04364 to REC). Two-photon characterization work (MD) was supported by the NIH BRAIN grant U24 NS109107 (Resource for Multiphoton Characterization of Genetically-Encoded Probes). Two-photon Ca^2+^ imaging of AP and all-optical data (IB and VE) were supported by the ERC Advanced (Grant ERC-2019-AdG; award no. 885090), the Medical Research foundation FRM (FRM-FDT202204015069) and the Axa research foundation. AA and REC wrote the manuscript. AA, SS, MD, LZ, JZ, S-YW, and YS designed and performed experiments and analyzed the data. IB and VE designed experiments for AP imaging under two-photon excitation and all-optical experiments. IB performed and analyzed the data of the corresponding the experiments. AWL, AGT, VE, KP and REC acquired funding and supervised the project. All authors contributed to editing and proofreading of the manuscript.

